# SPECTRA: predicting cellular perturbation responses with Graph Learning over Gene Regulatory Networks

**DOI:** 10.64898/2026.09.21.752624

**Authors:** Michele Calabrò, Patrick Sheehan, Francesco Cambuli, Andrea Sottoriva

## Abstract

Predicting the transcriptomic consequences of cellular perturbations is challenging due to the sparsity of single-cell transcriptional responses, heterogeneous perturbation efficiency, and the difficulty of identifying the small subset of genes that are truly differentially expressed after perturbation. Current approaches typically encode interventions as latent shifts from unperturbed to perturbed cellular state, making the models hard to interpret. Here we introduce SPEC-TRA — SPEctral CRISPR Transcriptome Regulatory Autoencoder — a graph-based model that treats CRISPR perturbations as interpretable, localized signals injected into a Gene Regulatory Network and propagated through directed graph neural networks. SPECTRA combines a variational control-cell encoder and a directed heterophilic graph decoder, integrating single-cell expression, pretrained transcriptomic context, and prior regulatory topology into a single node-level representation. We evaluate SPECTRA on a large-scale single-cell CRISPRi benchmark, measuring differentially expressed gene recovery through precision, F1 score, and AUPRC-based DEG classification. Since perturbations in SPECTRA are *interpretable*, we tested the predictions of the model against known biology, in cases where the effects of a gene knock out have been repeatedly demonstrated with orthogonal experiments. Our results show that graph-based signal propagation is a biologically grounded alternative to latent-shift perturbation modeling and can improve the recovery of sparse perturbation-induced transcriptional effects.

## 1 Introduction

One of the biggest goals of computational biology is the development of *virtual cell* models [1]; in its most comprehensive interpretation, this refers to computational models capable of simulating and reproducing the complex molecular mechanisms and interactions present in biological cells [2]. However, recently this term has increasingly been used to indicate data-driven Machine Learning models that predict the response state of a cell given a CRISPR or biochemical perturbation, with single-cell RNA-seq as a readout [3].

CRISPR-based technologies are particularly useful because they provide immediate insight into what happens inside a cell once a gene is perturbed. Perturbing a gene can reveal downstream transcriptional consequences, regulatory dependencies, and candidate therapeutic vulnerabilities [4]. However, because cellular responses depend highly on their specific biological context (e.g., cell type, disease state, or microenvironment), exhaustively testing all possible CRISPR perturbations across all genes and cell states *in vitro* is impractical due to the combinatorial explosion of possibilities [3]. Consequently, there is great interest in developing models capable of predicting the transcriptome of a cell in response to an unseen *in silico* perturbation, in order to dramatically narrow the biological search space and prioritize the most promising experiments.

Several models have been developed in recent years to tackle this challenge. A defining characteristic of these models is their shared foundational design: they learn a latent representation for the unperturbed cell state (the “basal state”) and combine it with a representation of the perturbation to learn a “shift” in the cell’s latent space. The post-perturbation gene expression profile is then generated via a “decoder” model that maps this shifted latent representation back into the high-dimensional gene space. While they share this common design of mapping control cells and shifting them via a perturbation embedding, these models differ in methodological flavour: CPA (Compositional Perturbation Autoencoder)[5] utilizes variational autoencoders (VAEs) coupled with adversarial training to disentangle the cell’s basal state from the perturbation embeddings, which are then recombined to predict unseen combinations; GEARS [6] uses graph neural networks (GNNs) over Gene Ontology (GO) graphs to learn the representation of the perturbation, and over gene co-expression networks to learn the representation of the unperturbed cell state; scLAMBDA [7] adopts a VAE approach that integrates gene embeddings derived from large language models (LLMs) to disentangle basal cell variations from perturbation-specific “salient” representations; STATE [8] employs a multi-scale Transformer architecture, uniquely utilizing self-attention mechanisms across sets of unperturbed control cells to capture underlying biological heterogeneity before predicting the perturbed state; MORPH [9] uses a discrepancy-based VAE with attention to model perturbation responses across modalities and contexts.

Despite rapid development, models still struggle with out-of-distribution (OOD) generalization to novel cellular contexts or unseen perturbations. Recent benchmarking reveals that complex models sometimes fail to outperform simple linear or mean-based baselines when tested on data distinct from their training sets [10]. To ensure that virtual cells truly capture generalizable biological logic rather than dataset-specific artifacts, the field is increasingly relying on standardized evaluation frameworks. The Virtual Cell Challenge of 2025 [3] framed this task as a community benchmark for predicting cellular response to perturbations and provides purpose-built datasets and evaluation criteria for measuring progress toward this goal.

We maintain that the primary practical utility of “virtual cell” models in genetic perturbation settings is not merely to reconstruct global transcriptomes, but to identify the often sparse set of genes and pathways that are specifically altered by an intervention, in order to learn the underlying biology explicitly. This motivates the design of a model that is structurally biased toward sparse regulatory propagation rather than global latent interpolation. Here we present SPECTRA (SPEctral CRISPR Transcriptome Regulatory Autoencoder) a model that addresses this need by representing a CRISPR perturbation as a localized graph signal injected at the perturbed gene and propagated through a directed Gene Regulatory Network. In contrast to latent-shift models, SPECTRA explicitly constrains perturbation effects to propagate through biologically plausible regulatory topology while learning to re-weight noisy edges through directed, heterophily-aware graph convolution, and accounts for the heterogeneity and unpaired nature of the data using a variational approach.

To rigorously assess the predictive capabilities of SPECTRA, we employed a dual approach: a) test the model predictions against large Pertub-seq dataset and b) ensure the model predicts the downstream effects of a set of well characterized perturbations observed with orthogonal experiments in many model systems. Here we show that SPECTRA achieves state-of-the-art results, demonstrating that the strategy of propagating perturbation signals across a biological knowledge graph provides a highly effective and robust alternative to the conventional “latent shift” paradigm widely adopted by current perturbation models.

## 2 Results

### 2.1 Model Overview

SPECTRA is a graph-based variational autoencoder designed to predict transcriptomic responses to CRISPR perturbations by explicitly modeling perturbations as signals propagating through a directed Gene Regulatory Network (GRN), rather than as global shifts in the latent state of a cell (Figure 1a).

**Figure 1:**
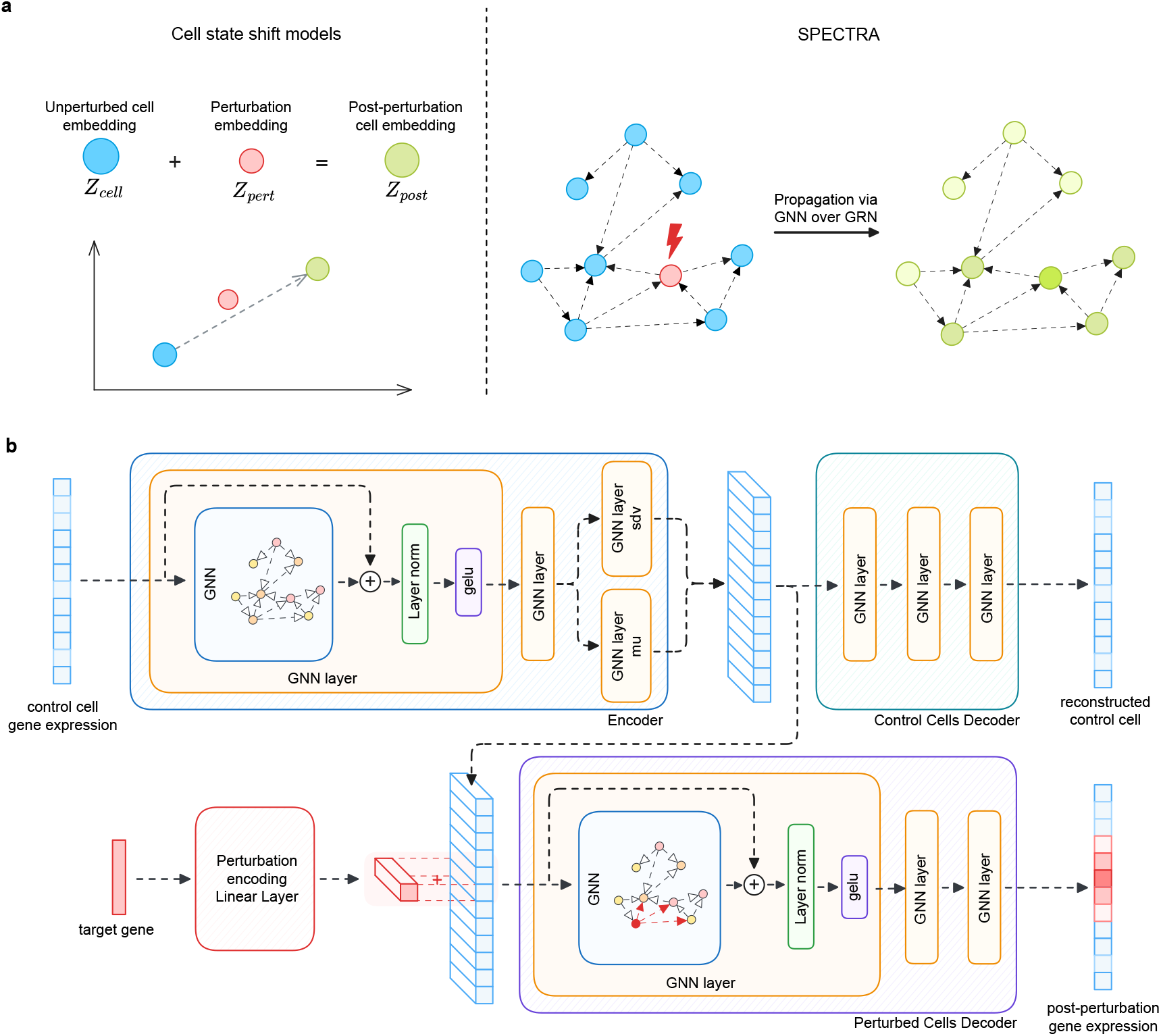
Overview of the SPECTRA framework. (a) Conceptual paradigm. A comparison of SPECTRA (SPectral CRISPR Transcriptome Regulatory Autoencoder) with conventional virtual cell architectures. While standard models predominantly represent genetic interventions as global shifts within a cell’s latent state space, SPECTRA explicitly models perturbations as localized signals propagating through a Gene Regulatory Network (GRN) utilizing graph neural networks. (b) Model architecture. A Variational Graph Autoencoder (VGAE) framework is employed to learn a regularized representation of the unperturbed (basal) cell state. Concurrently, a perturbation encoder generates a latent embedding for the intervened gene. This embedding is directly integrated with the target gene’s node-specific representation within the basal state, establishing a structurally localized perturbation signal. A GNN decoder then simulates the resulting downstream regulatory cascade by propagating this localized signal across the directed GRN topology to predict the final post-perturbation transcriptomic profile.

The architecture consists of three main components: a graph encoder that learns a latent representation of the basal cellular state, a perturbation encoder that generates a latent representation of the targeted gene, and a graph decoder that simulates the downstream propagation of the perturbation across the regulatory network.

The model receives as input the gene expression profile of an unperturbed control cell, combined with pretrained gene embeddings obtained from the scGPT foundation model [11]. These embeddings are conditioned on the observed expression of each gene through a Feature-wise Linear Modulation (FiLM) layer [12], allowing each node representation to integrate cell-specific transcriptional information together with the broader biological context encoded by the pretrained embeddings. The resulting node features are processed by a variational graph encoder to construct a graph-structured latent representation of the basal cell state while simultaneously regularizing the latent space through a Graph Variational Autoencoder objective [13].

Unlike conventional perturbation models, which typically combine a perturbation embedding with the entire latent representation of the cell, SPECTRA injects the perturbation only into the latent representation of the targeted gene. The perturbed latent graph is then decoded through multiple graph convolution layers that learn how this localized signal propagates through the GRN to produce downstream transcriptional changes. To accurately model the topology of regulatory networks, the decoder employs a directed adaptation of Frequency Adaptation Graph Convolution Networks (Dir-FAGCN) [14, 15]. This architecture explicitly accounts for both graph directionality and heterophily while learning edge-specific propagation coefficients, allowing the model to emphasize biologically relevant regulatory interactions and attenuate noisy or spurious connections. Together, these design choices enable SPECTRA to combine prior biological knowledge, pretrained transcriptomic representations, and graph-based signal propagation into a unified framework for perturbation prediction.

### 2.2 SPECTRA improves recovery of perturbation-specific transcriptional responses

We trained and evaluated SPECTRA on the Virtual Cell Challenge dataset [3]. This dataset comprises approximately 300,000 single-cell RNA-seq profiles from H1 human embryonic stem cells (hESCs) following CRISPR interference (CRISPRi) silencing of 150 target genes.

To assess the ability of SPECTRA to generalize to unseen genetic perturbations, we split the dataset at the perturbation level rather than at the cell level. In this setting, all cells associated with a given perturbation are assigned exclusively to either the training, validation, or test set, ensuring that perturbations evaluated at validation and test time are never observed during training. We used 75% of perturbations for training, 15% for validation, and 15% for testing. This split defines a perturbation-generalization task [3], in which the model must predict the post-perturbation transcriptional response of CRISPRi interventions not seen during training.

We evaluated SPECTRA using a comprehensive set of metrics designed to capture both global transcriptomic similarity and perturbation-specific biological signal. Following recent benchmarking frameworks [16], we report population-average metrics, including Mean Squared Error (MSE), PCC-delta, and E-distance [17], as well as distribution metrics, including Wasserstein distance and KL divergence. Because genetic perturbations often induce sparse transcriptional responses, we compute these metrics both across all genes and on a finite set of perturbation-specific Differentially Expressed Genes (DEGs) for a more focused evaluation. In addition, we explicitly evaluate DEG recovery by treating each perturbation as a binary classification task over genes and reporting F1 score and AUPRC [18].

We compare SPECTRA against established perturbation prediction models, including GEARS [6] and scLAMBDA [7], as well as a non-parametric pseudobulk baseline. Full details on preprocessing, model training, baseline implementation, and metric computation are provided in the Methods.

Across 10 independent runs per model, SPECTRA showed a distinct performance profile compared with the other models (Figure 2a). On unweighted MSE, SPECTRA did not outperform the pseudobulk baseline or the alternative deep learning models. Specifically, SPECTRA obtained an MSE almost reaching 0.0044, whereas GEARS, the pseudobulk baseline, and scLAMBDA achieved values below 0.003. This result is consistent with the known limitation of global reconstruction metrics in sparse perturbation settings: because most genes remain unchanged after CRISPRi perturbations, models that predict near-average expression profiles can achieve low global error without accurately recovering the subset of genes that truly respond to the perturbation [19].

**Figure 2:**
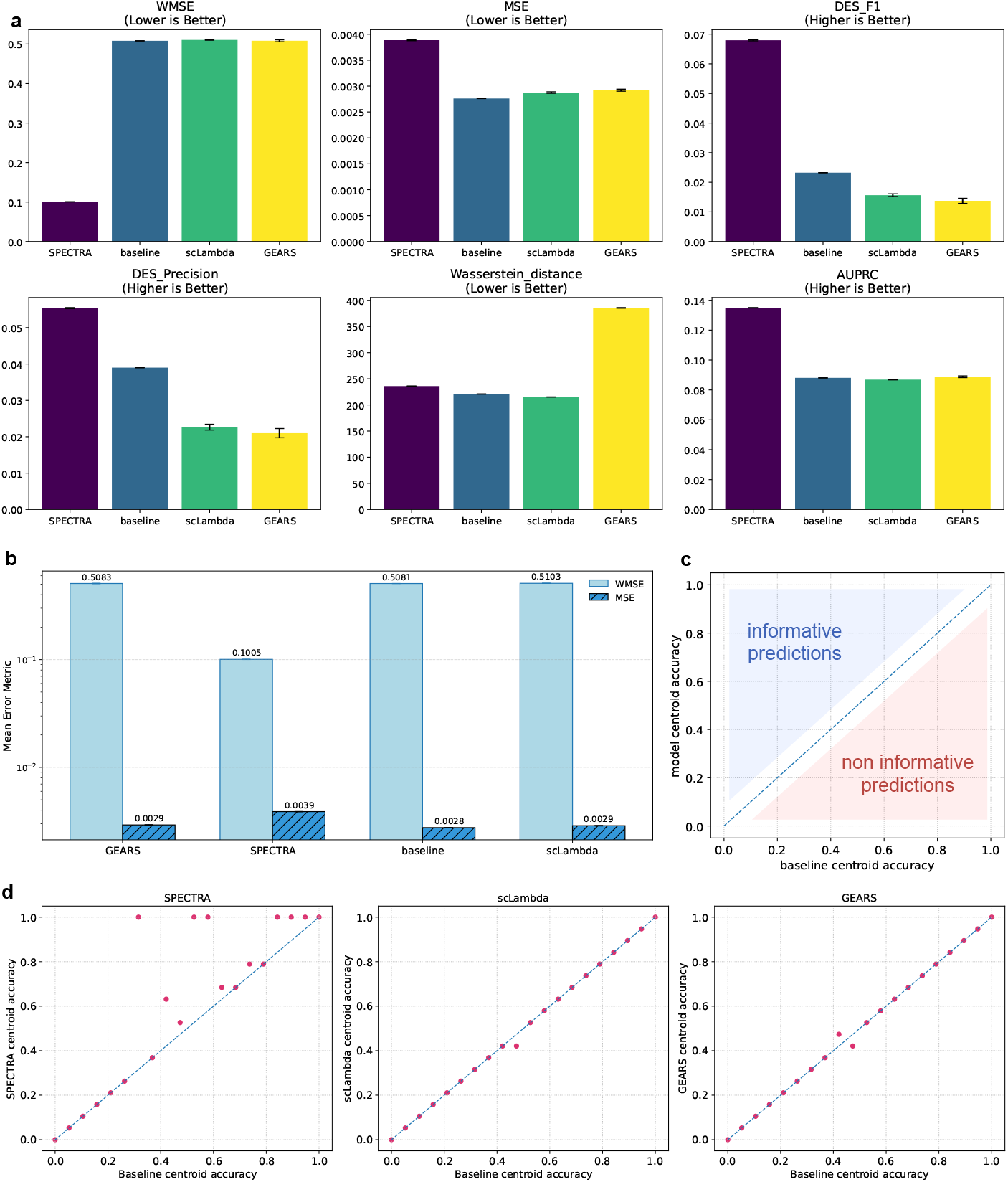
Benchmarking SPECTRA against GEARS, scLAMBDA, and a pseudobulk baseline on held-out Virtual Cell Challenge CRISPRi perturbations, averaged over 10 runs per model. (a) Comparison of global reconstruction, distributional, and differentialexpression metrics. Notably, SPECTRA achieves the highest performance on DEG-overlap metrics. (b) Comparison of MSE and DEG-weighted MSE as defined in [19]. SPECTRA achieves substantially lower WMSE than all evaluated alternatives, despite slightly higher unweighted MSE. (c) Schematic interpretation of centroid-accuracy comparisons as defined in the Systema framework [20]: points above the diagonal correspond to model predictions that improve over the pseudobulk baseline, whereas points near or below the diagonal indicate baseline-like or non-informative predictions. (d) Perturbation-level centroid accuracy for SPECTRA, scLAMBDA, and GEARS compared with the pseudobulk baseline; SPECTRA substantially improves the recovery of perturbation-specific effects compared with the other models.

By contrast, when errors were weighted toward perturbation-responsive genes, SPECTRA sub-stantially outperformed all evaluated alternatives. SPECTRA achieved an average WMSE of 0.1005 across 10 runs, compared with 0.5083 for GEARS, 0.5081 for the pseudobulk baseline, and 0.5103 for scLAMBDA (Figure 2b). Thus, although SPECTRA does not minimize global transcriptome-wide reconstruction error, it is the most precise model in accurately capturing the expression changes concentrated in genes with stronger perturbation-associated differential expression.

This improvement was also reflected in DEG recovery metrics. Treating differential-expression prediction as a binary classification task over genes, SPECTRA achieved the highest differential-expression F1 score, AUPRC and precision among all evaluated models (Figure 2a). These results indicate that SPECTRA is better able to identify the genes whose expression changes after perturbation, which is the primary biological use case of in silico CRISPR response prediction.

Distributional metrics showed a more nuanced pattern. SPECTRA achieved a Wasserstein distance comparable to the pseudobulk and scLAMBDA baselines and lower than GEARS, but it did not uniformly dominate all distribution-level criteria. Taken together, these results suggest that the main advantage of SPECTRA is not simply improved global reconstruction of the post-perturbation transcriptome, but improved recovery of the sparse, perturbation-specific component of the response.

Finally, we evaluated whether each model learned perturbation-specific effects or merely reproduced the average perturbation response. To this end, we adopted the centroid-accuracy analysis proposed by the Systema framework [20], comparing each model against the non-parametric pseudobulk baseline. Centroid accuracy is a metric that measures whether predicted post-perturbation profiles are closer to their correct ground-truth centroid than to the centroids of other perturbations. In our representation, predictions along and below the diagonal are effectively uninformative, as they are no closer to the correct perturbation centroid than the baseline, which captures only the average perturbed state. Conversely, predictions above the diagonal indicate recovery of perturbation-specific effects beyond this shared systematic component (Figure 2c). Under this criterion, SPECTRA was the only evaluated model showing a clear improvement over the baseline for a subset of held-out perturbations (Figure 2d), whereas GEARS and scLAMBDA largely remained aligned with the diagonal. These results suggest that, unlike the competing models, SPECTRA can recover perturbation-specific transcriptional structure that is not explained solely by systematic differences between control and perturbed cells.

### 2.3 Benchmarking on Standardized Virtual Cell Evaluation Framework

To contextualize the performance of SPECTRA against standard community benchmarks, we evaluated our predictions on the held-out test perturbations using the vcc eval metric suite [8] alongside our established evaluation criteria (Figure 3).

**Figure 3:**
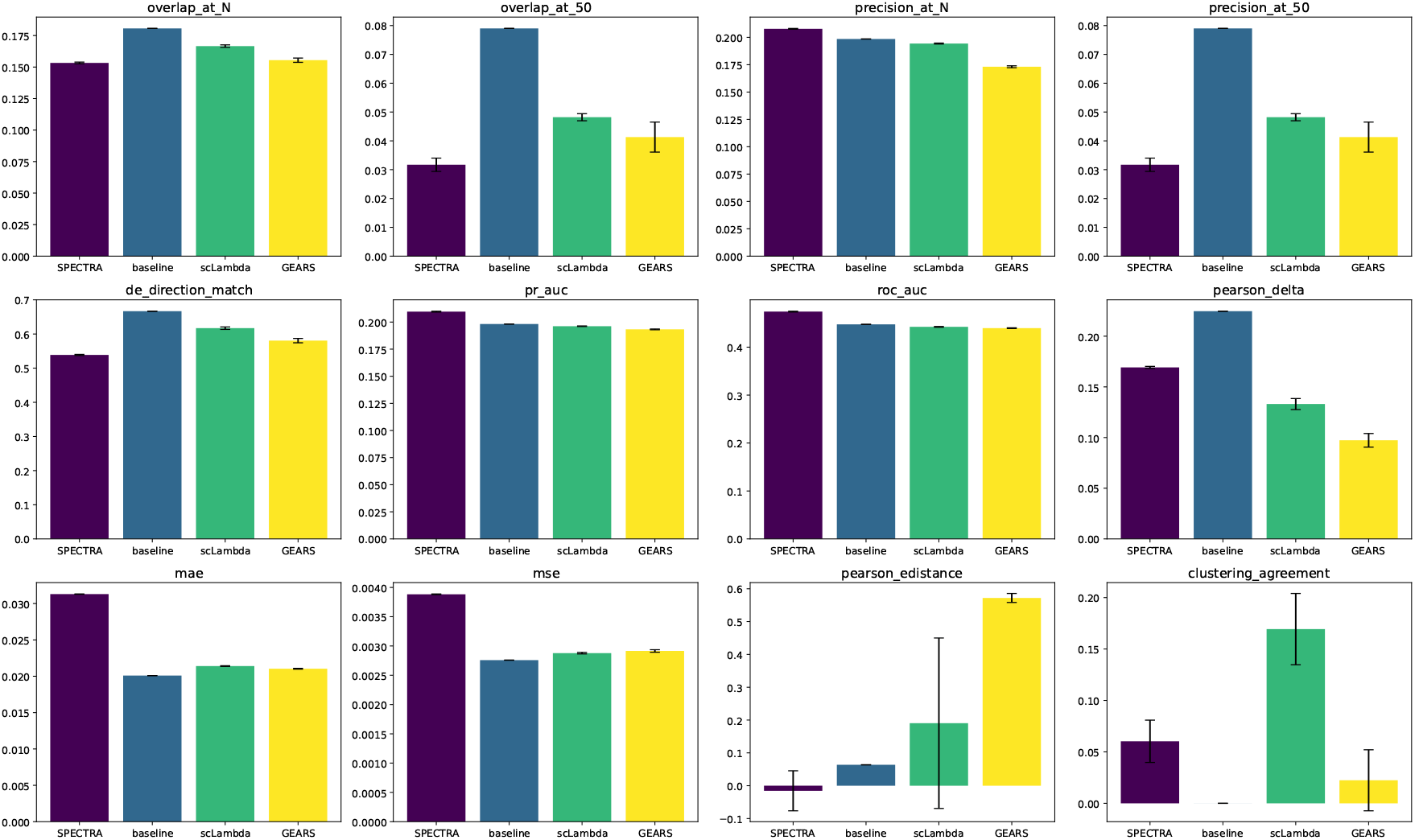
Benchmarking SPECTRA against GEARS, scLAMBDA, and a pseudobulk baseline on held-out Virtual Cell Challenge CRISPRi perturbations using vcc eval suite.

SPECTRA exhibits a distinct trade-off across the evaluated metrics. On summary error metrics such as Mean Absolute Error (MAE), SPECTRA does not uniformly outperform mean-based baselines or unconstrained generative models. As explained above, this behavior is expected: global point-wise distances are overwhelmingly dominated by the vast majority of unaffected, non-differentially expressed genes, which rewards conservative predictors that collapse toward the global average expression profile [19]. Similarly, Pearson correlation is known to be confounded by systematic variation—the shared background transcriptional shift induced across all perturbation conditions—enabling trivial predictors to achieve strong correlation scores without resolving target-specific regulatory responses [20].

In contrast, SPECTRA demonstrates superior performance on classification and discrimination metrics that directly evaluate DEG recovery, achieving the highest area under the precision-recall curve (PR-AUC = 0.210) and receiver operating characteristic curve (ROC-AUC = 0.472) among all benchmarked architectures, including GEARS and scLAMBDA.

While standard vcc eval formulations evaluate DEG identification via PR-AUC, they define true-positive labels through a fixed statistical significance threshold (e.g., Wilcoxon *p*_adj_ *<* 0.05). This hard-thresholding approach introduces boundary artifacts, where genes with biologically meaningful fold changes or borderline significance are arbitrarily penalized. In our primary evaluation framework, we compute an effect-size-modulated AUPRC by continuously evaluating recovery across varying effect-size (log FC) thresholds [18]. This formulation avoids binarization artifacts and directly assesses the model’s ability to rank true biological effect sizes rather than statistical test sensitivity alone.

Furthermore, we intentionally de-emphasize fixed overlap metrics such as Overlap@*N*. In standard benchmarks, Overlap@*N* effectively operates as a recall metric evaluated at the true DEG set size *N*. While recall measures sensitivity to true positives, it fails to penalize false positives—a critical drawback in single-cell perturbation screening where candidate validation requires costly follow-up wet-lab experiments. In this regime, false discoveries carry a high experimental cost. Consequently, our primary benchmarking prioritizes precision and *F*_1_ score, ensuring that predicted regulatory cascades reflect high-confidence, perturbation-specific transcriptional consequences.

## 3 SPECTRA propagation paths recover target-specific response programs in hESCs

SPECTRA enables the extraction and interpretation of the signed edge weights produced by the FAGCN layers of the decoder. Unlike standard attention scores, FAGCN coefficients can be interpreted as homophily-aware propagation weights [14], with positive coefficients favoring smoothing and low-frequency propagation between connected nodes, and negative coefficients favoring feature differentiation and high-frequency propagation, thereby allowing the model to represent heterophilic relationships. Importantly, the sign of a FAGCN coefficient is not necessarily equivalent to transcriptional activation or repression; the direction of the predicted expression response was therefore assessed separately using pseudobulk log_2_ fold changes.

To summarize propagation across successive graph-convolution layers, we defined a multiplicative score for each path originating from the perturbed gene and spanning one to three hops:

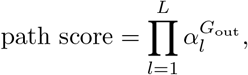

where 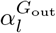 denotes the FAGCN gate weight associated with edge *l*, considering only the down-stream direction defined by *G*_out_ (see Methods). For each path length *L* ∈ {1, 2, 3}, we retained the *K* = 20 highest-scoring paths, ranked according to the absolute value of their path score. Importantly, these paths represent layer-wise routes of message propagation rather than temporal molecular sequences. Moreover, because the path score considers only the FAGCN coefficients and does not account for feature transformations, residual or initial-state contributions, nonlinearities, or subsequent decoder operations, it should be interpreted as a heuristic summary of network reweighting rather than as an exact attribution of the final prediction. In addition, SPECTRA can only reweight interactions already present in the prior graph and therefore cannot recover missing edges.

We qualitatively examined three curated targets selected *a priori* because they are associated with distinct and well-characterized responses in human embryonic stem cells: broad loss of pluripotency and multilineage differentiation (POU5F1), neuroectoderm-biased differentiation (NANOG), and p53-mediated stress, cell-cycle arrest, and apoptosis (MDM2). The aim of this analysis was to assess whether the downstream network reweighting learned by SPECTRA is consistent with these established biological responses. SPECTRA was trained on the H1 hESC CRISPR-interference dataset generated for the Virtual Cell Challenge; consequently, its predicted responses may be weaker than, or differ from, phenotypes reported after complete Cas9 knockout or other types of experiment [3, 21].

For each perturbation, we generated 256 cells and averaged the corresponding FAGCN edge weights across the predicted cell population. Predicted pseudobulk log_2_ fold changes were then calculated relative to non-targeting controls for genes contained within the selected downstream subgraphs.

### 3.1 POU5F1 silencing predicts broad exit from pluripotency

POU5F1 encodes OCT4, a core regulator of pluripotency. In primed human pluripotent stem cells, POU5F1 knockout produces a strong exit from pluripotency and heterogeneous differentiation toward both neuroectodermal and mesendodermal states [21]. Consistent with this broad phenotype, SPECTRA identifies a collection of genes that look consistent with exit from pluripotency and acquisition of differentiated developmental programs, including evidence for more than one lineage. Particularly informative candidate routes include POU5F1 → USP9X → CER1 and POU5F1 → DCLK1 → PROX1 → NTN1, in addition to other paths terminating in NFIA, DCX, KALRN, and MYOCD (Figure 4a), collectively indicating several different lineage-associated outcomes. At the expression level (Figure 4b), LIN28A, a marker and functional regulator of the undifferentiated hPSC state, as well as a direct OCT4 target [22], is among the most strongly downregulated genes in the cascade [23, 24]. Conversely, CER1 and NOTO are predicted to be upregulated, consistent with primitive-streak/mesendodermal and axial-mesodermal programs, respectively. Several neural- or neurodevelopment-associated genes, including NFIA, NTN1, PROX1, DCX, KALRN, DCLK1, KCNQ5, KCNMA1, and ADAM22, are also predicted to increase. The coexistence of neural and mesendodermal endpoints is therefore compatible with the multilineage response reported after POU5F1 loss. However, beyond LIN28A, canonical pluripotency factors such as NANOG, SOX2, PRDM14, and SALL4 are absent from the displayed top paths. Furthermore, many intermediate paths found by SPECTRA are noncanonical and therefore require independent validation before being considered as true molecular causal pathways. Because of this, our analysis supports convergence on an appropriate differentiated state more strongly than it supports complete reconstruction of the canonical OCT4 regulatory causal chain.

**Figure 4:**
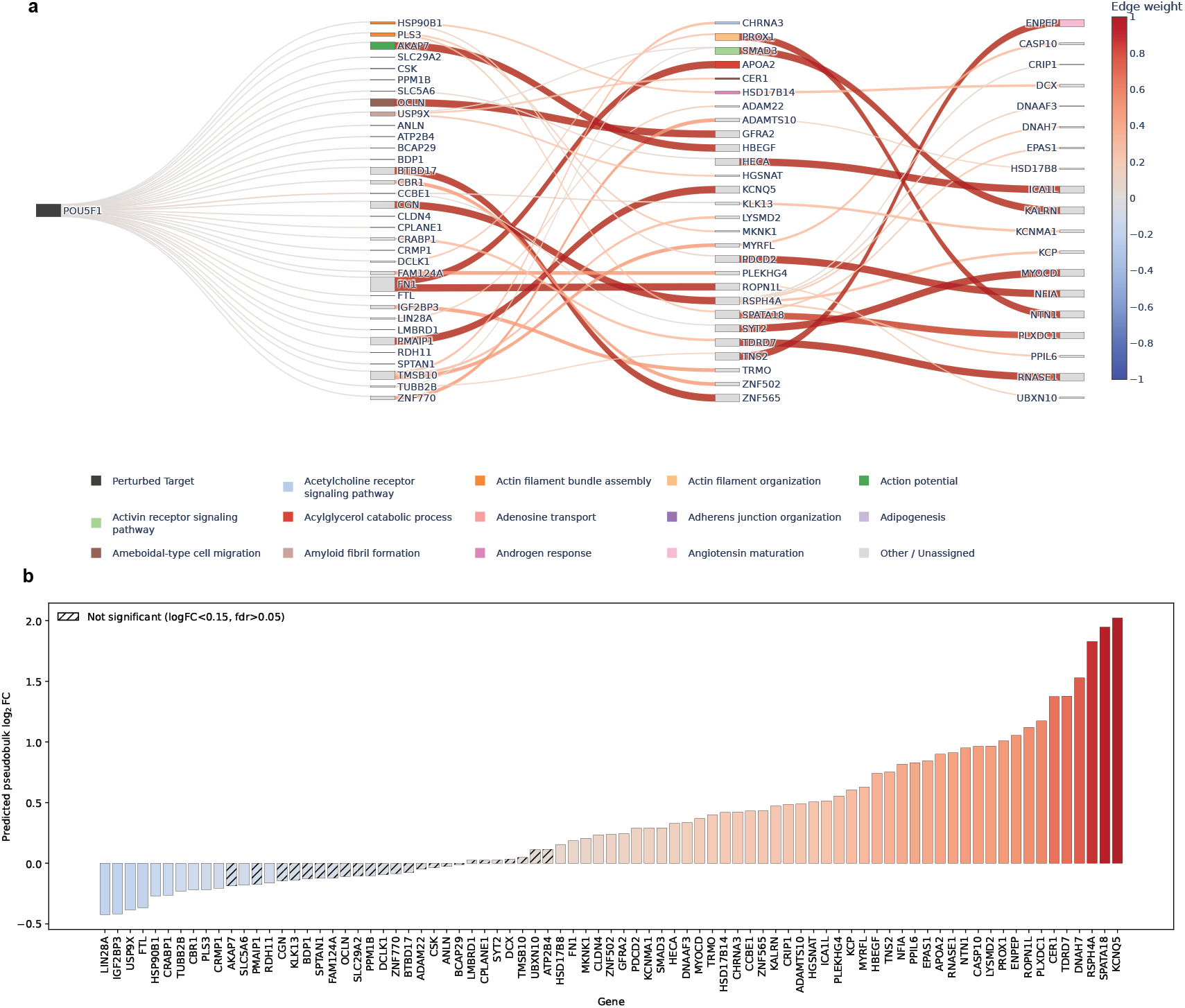
SPECTRA propagation analysis following POU5F1 *in silico* silencing. (a) Sankey representation of the union of the top *K* = 20 one-, two-, and three-hop paths originating from POU5F1, ranked within each path length according to the absolute value of the multiplicative path score. Edges are colored according to their signed FAGCN coefficients, while node annotations are derived from functional enrichment analysis performed with `gseapy` using `MSigDB_Hallmark_2020` and `GO_Biological_Process_2026`. (b) Predicted pseudobulk log_2_ fold changes relative to non-targeting ground-truth controls for genes contained in the selected down-stream subgraph. Genes with log_2_ FC *<* 0.15 or FDR-adjusted *p >* 0.05 are marked as non-significant.

### 3.2 NANOG silencing predicts neuroectoderm-biased differentiation

NANOG loss also destabilizes pluripotency, but in primed hPSCs the response is more lineage-biased, with preferential differentiation toward neuroectoderm rather than the broader mixture observed after POU5F1 perturbation [21]. SPECTRA prioritizes several distinct paths converging on neurodevelopment-associated genes (Figure 5a), including NANOG → IRX2 → IRX1, NANOG → TERF1 → FOXB1 → ASTN1, NANOG → ZBTB3 → CAMK2D → SORCS3, and NANOG → TERF1 → CAMK2D → SORCS3 (Figure 5). MAP2 and IRX1 are among the most strongly upregulated genes in the selected cascade; notably, MAP2 is also used as a neural marker in the independent hPSC Perturb-seq analysis [[21], Fig. S3]. Additional selected genes, including IRX2, FOXB1, SEMA6A, ASTN1, GRIA3, GRIK3, SORCS3, and RORA, further support a neural-developmental endpoint. The one-hop neighbourhood also contains developmental-signalling components such as BMPR1A, associated with BMP signaling which is intimately involved in hESC fate determination, and SFRP1, together with the developmental transcription factor SOX4. Nevertheless, the selected cascade does not show a clear coordinated decrease in POU5F1, PRDM14, or SALL4. As in the POU5F1 case, SPECTRA therefore captures the lineage-acquisition component more clearly than the collapse of the core pluripotency network.

**Figure 5:**
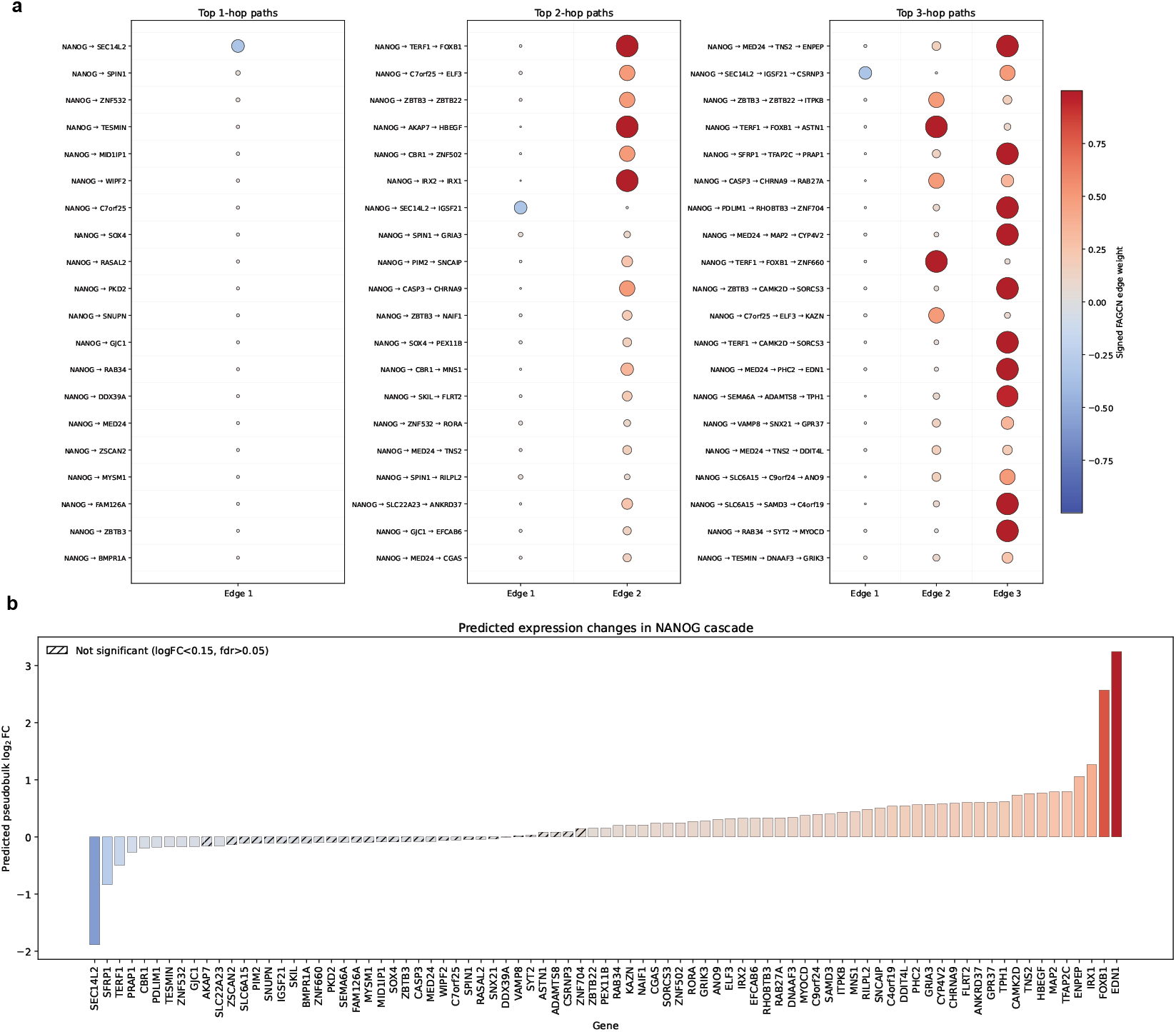
SPECTRA propagation analysis following NANOG silencing. (a) Top *K* = 20 one-, two-, and three-hop paths originating from NANOG, ranked within each path length according to the absolute value of the multiplicative path score. Each row represents a selected path and columns correspond to successive edges along the path. Dot color represents the signed FAGCN coefficient, while dot size reflects its absolute magnitude. (b) Predicted pseudobulk log_2_ fold changes relative to non-targeting controls for genes contained in the selected downstream paths.

### 3.3 MDM2 silencing predicts activation of a p53-centred stress program

MDM2 encodes a major negative regulator of p53; its loss stabilizes p53 and can elicit cell-cycle arrest, stress responses, and apoptosis [25, 26]. In H1 hPSCs, MDM2-targeting guides are depleted in genome-wide CRISPR screens, consistent with reduced cellular fitness following MDM2 loss [27]. SPECTRA recovers a particularly coherent one-hop neighbourhood containing several established p53-responsive genes (Figure 6a), including CDKN1A, a canonical p53-mediated cell-cycle arrest effector, BTG2, TP53INP1, GDF15, DRAM1, GLS2, TNFRSF10A, PLK3, and MDM4. As MDM2 perturbation is simulated in the context of H1 embryonic stem cells, it is also interesting to observe a strong interaction with the developmental node made up by NODAL and BMP4, two key regulators of pluripotency differentiation, previously linked to MDM2 function [28]. Predicted pseudobulk changes (Figure 6b) further show increases in CDKN1A, BTG2, DRAM1, FAS, and GDF15, supporting activation of a p53-associated arrest and programmed cell death.

**Figure 6:**
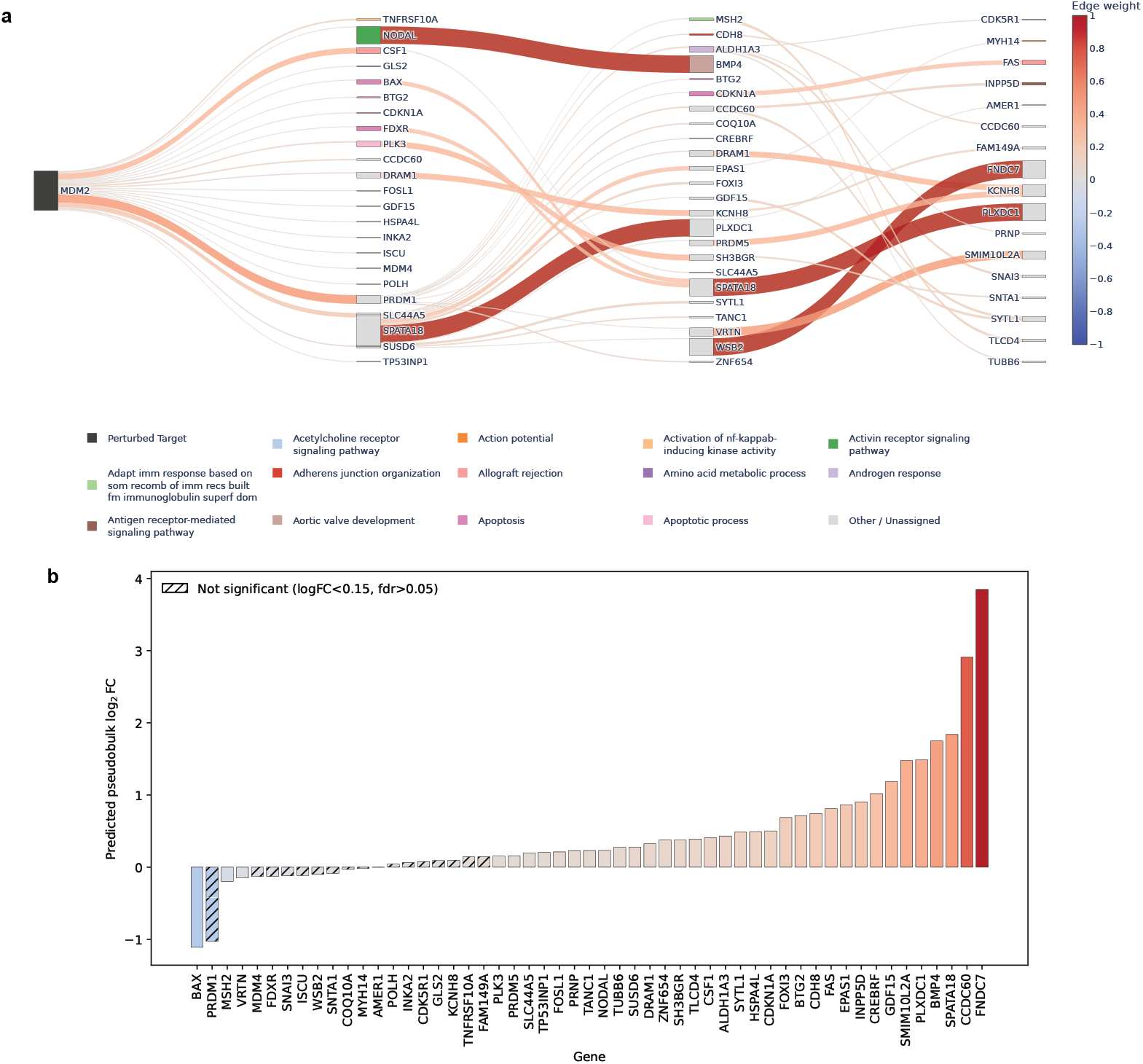
SPECTRA propagation analysis following MDM2 silencing. (a) Sankey representation of the union of the top *K* = 20 one-, two-, and three-hop paths originating from MDM2, ranked within each path length according to the absolute value of the multiplicative path score. (b) Predicted pseudobulk log_2_ fold changes relative to non-targeting controls for genes contained in the selected downstream subgraph.

Among the longer routes, MDM2 → PRDM1 → CDKN1A → FAS link antiproliferative and death-receptor components, whereas routes containing SPATA18 and DRAM1 are consistent with mitochondrial quality-control and autophagy-associated stress responses [29]. However, these exact chains are not necessarily established sequential mechanisms. For example, the selected MDM2 → BAX → SPATA18 → PLXDC1 path should not be interpreted literally, because BAX is primarily an apoptotic effector rather than an obvious transcriptional relay [30]. BAX is also predicted to be downregulated, in contrast to the simplest expectation under p53 activation. This discrepancy may reflect model error, incomplete CRISPRi, context- or time-dependent transcriptional responses, or the importance of post-transcriptional BAX regulation.

### 3.4 Cross-target interpretation

Across the three perturbations, the dominant endpoint programs are target-specific: POU5F1 paths involve neural and mesendodermal outcomes, NANOG paths converge more selectively on neurodevelopmental programs, and MDM2 paths recover a p53-centred stress, cell-cycle arrest, and apoptosis program. This cross-target specificity constitutes the strongest qualitative result of the analysis.

We further investigated whether the signed FAGCN coefficients were associated with gene–gene coexpression patterns in the predicted perturbed state. For each perturbation, Pearson and Spearman correlations were calculated between the FAGCN edge weights and the corresponding gene–gene Pearson expression correlations across the 256 generated perturbed cells, considering only edges belonging to the top-ranked paths described above. Signed FAGCN coefficients were positively and significantly correlated with gene–gene coexpression for POU5F1 and NANOG, but not for MDM2 (Table 1). We extended this analysis to a selection of additional perturbations, including genes associated with activation of the p53 pathway, and observed a similar positive association for most of the examined targets (Table 1).

**Table 1:** Association between signed FAGCN edge weights and gene–gene coexpression across perturbation-specific propagation paths. For each target gene, gene–gene coexpression was calculated across 256 SPECTRA-predicted perturbed cells for edges contained in the selected top-ranked paths. Pearson and Spearman correlation coefficients quantify the association between the signed FAGCN edge weights and the corresponding gene–gene coexpression values. Reported *p*-values test the null hypothesis of zero correlation; the final column indicates the number of edges included in each analysis.

| target gene | pearson r | pearson $p$-value | spearman r | spearman $p$-value | number of edges |
| --- | --- | --- | --- | --- | --- |
| NUPR1 | 0.77 | $1.77 \times 10^{-10}$ | 0.42 | $3.25 \times 10^{-3}$ | 48 |
| PMAIP1 | 0.62 | $4.10 \times 10^{-10}$ | 0.53 | $2.67 \times 10^{-7}$ | 83 |
| PRAP1 | 0.58 | $8.54 \times 10^{-6}$ | 0.54 | $4.47 \times 10^{-5}$ | 51 |
| SESN2 | 0.46 | $1.56 \times 10^{-5}$ | 0.28 | $1.24 \times 10^{-2}$ | 80 |
| TRIAP1 | 0.43 | $4.83 \times 10^{-5}$ | 0.34 | $1.77 \times 10^{-3}$ | 82 |
| BAX | 0.34 | $1.04 \times 10^{-3}$ | 0.24 | $2.68 \times 10^{-2}$ | 88 |
| PLK3 | 0.40 | $1.49 \times 10^{-3}$ | 0.07 | $5.88 \times 10^{-1}$ | 61 |
| PLK2 | 0.28 | $1.27 \times 10^{-2}$ | 0.21 | $6.63 \times 10^{-2}$ | 76 |
| PHLDA3 | -0.25 | $2.45 \times 10^{-2}$ | -0.35 | $1.63 \times 10^{-3}$ | 79 |
| DDIT4 | 0.22 | $5.90 \times 10^{-2}$ | 0.14 | $2.23 \times 10^{-1}$ | 76 |
| CDKN1A | -0.10 | $4.12 \times 10^{-1}$ | -0.26 | $2.76 \times 10^{-2}$ | 71 |
| MDM2 | 0.06 | $6.23 \times 10^{-1}$ | 0.02 | $8.39 \times 10^{-1}$ | 69 |

This result is consistent with the interpretation of positive FAGCN coefficients as favouring concordant signals and negative coefficients as favouring divergent signals [14]. More generally, the observed association indicates that the edge weights learned by SPECTRA are related to measurable biological structure in the predicted expression profiles, rather than representing arbitrary model parameters. However, this relationship is not universal, as for example illustrated by MDM2 and some other genes in Table 1, and FAGCN coefficients should therefore not be interpreted simply as measures of gene coexpression. Instead, this correlation analysis provides complementary evidence that, for several perturbations, the network reweighting learned by SPECTRA captures biologically meaningful relationships between connected genes.

## 4 Discussion

Current virtual cell models face several fundamental challenges. Most notably, the destructive nature of single-cell RNA sequencing prevents paired observations of the same cell before and after perturbation, forcing models to infer perturbation effects from unpaired population-level distributions affected by technical noise and biological heterogeneity. Additional sources of variability arise from incomplete perturbation efficacy, heterogeneous knockdown efficiency in CRISPRi experiments, off-target sgRNA activity, and intrinsic transcriptional programs, such as cell-cycle progression, which can obscure the subtle and sparse gene expression changes induced by genetic perturbations.

Another major challenge lies in model evaluation. Conventional metrics such as MSE (Mean Square Error), MAE (Mean Average Error), Pearson correlation, and *R*^2^ primarily assess global transcriptomic similarity and can therefore reward models that predict near-mean expression, despite failing to recover the sparse, perturbation-specific transcriptional changes induced by CRISPR interventions [19, 20, 31]. Since the primary biological objective of virtual cell models is to identify the genes that truly respond to perturbation, evaluation should also emphasize Differentially Expressed Gene (DEG) recovery rather than global expression reconstruction.

In this context, SPECTRA introduces a graph signal propagation framework for predicting CRISPR-induced transcriptomic responses. Instead of representing perturbations as global shifts in latent cell-state space, SPECTRA injects a local signal at the perturbed gene and learns to propagate that signal through a directed Gene Regulatory Network. This architectural choice is motivated by two biological observations: CRISPR perturbations often produce sparse transcriptional responses, and regulatory interactions among genes are themselves sparse and directional.

The model is designed to address several limitations of current virtual cell models. First, by operating directly over a GRN, SPECTRA incorporates prior biological structure and constrains the space of possible perturbation effects. Second, by using heterophily-aware FAGCN layers, SPECTRA avoids the inappropriate assumption that connected genes must have similar expression profiles. Third, by separating incoming and outgoing regulatory messages, SPECTRA respects the directionality of regulatory cascades. Fourth, by using DEG-weighted loss terms, the model explicitly prioritizes sparse perturbation-induced shifts rather than global transcriptome similarity.

SPECTRA’s evaluation philosophy follows recent critiques of perturbation-model benchmarking. Global metrics remain useful for assessing whether the model produces plausible expression profiles, but they are insufficient to determine whether the model captures biologically actionable perturbation effects. DEG recovery is a more direct proxy for how virtual cells are likely to be used in practice: prioritizing genes, pathways, and perturbations for experimental validation. The AUPRC metric is particularly appropriate because it measures the ranking of true DEGs above non-DEGs and is robust to threshold choices.

SPECTRA also offers an interpretability advantage. Because perturbations propagate through explicit graph edges, the learned FAGCN propagation weights can be inspected to identify candidate regulatory paths. This distinguishes SPECTRA from models in which predictions are generated from dense latent representations that are difficult to map back to gene-gene regulatory mechanisms.

The main limitation of SPECTRA is its dependence on GRN quality. Inferred GRNs can contain false-positive edges, missing interactions, context-inappropriate regulatory links, and ambiguous directionality [32]. Future versions of SPECTRA should therefore explore dynamic or perturbation-adaptive graph refinement, ensemble GRNs, confidence-weighted edges, and graph Transformer layers that can attend beyond the fixed local neighborhood.

In conclusion, SPECTRA reframes genetic perturbation prediction as a graph signal propagation problem over directed Gene Regulatory Networks. By combining expression-conditioned gene embeddings, variational control-cell encoding, localized perturbation injection, directed heterophilic graph propagation, and DEG-weighted distributional training, SPECTRA provides a biologically grounded alternative to latent-shift virtual cell models. Its focus on DEG recovery directly addresses the main practical use case of in silico perturbation modeling: identifying genes and pathways that are likely to change after intervention. Final quantitative benchmarking will determine the magnitude of SPECTRA’s advantage, but the model’s design establishes a clear and interpretable framework for sparse, network-aware perturbation prediction.

## 5 Data and Code Availability

The single-cell RNA-sequencing data from the Virtual Cell Challenge used to train and evaluate the model are publicly available at www.virtualcellchallenge.org. The source code, custom algorithms, and scripts required to reproduce the analyses presented in this manuscript are openly available on GitHub at www.github.com/MCalabroCode/SPECTRA.

## 6 Acknowledgments

This work was supported by the Associazione Italiana per la Ricerca contro il Cancro (AIRC) to A.S. (28961). A.S. is also supported by the ERC Consolidator Award (101125077). We thank Konstantin Winter and Anastasiia Romanova for the invaluable support to the Computational Biology Research Centre at Human Technopole.

## Methods

### Problem formulation

Let *G* = (*V, E*) denote a directed Gene Regulatory Network, where each node *v* ∈ *V* corresponds to a gene and each directed edge (*u, v*) ∈ *E* represents a putative regulatory relationship from gene *u* to gene *v*. Let 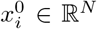 with *N* = |*V* | denote the log-normalized expression profile of an unperturbed control cell *i*, and 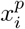 the log-normalized expression profile of cell *i* after perturbing gene *g*_*p*_. With a slight abuse of notation, we will often use *x* instead of *x*_*i*_ to indicate a single cell gene expression profile. The goal of SPECTRA is to predict samples from the post-perturbation expression distribution *P* (*X*^*p*^), given control cells sampled from *P* (*X*^0^), the perturbation target *g*_*p*_, and the regulatory graph *G*.

Because scRNA-seq is destructive, the model never observes paired samples 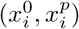 for the same biological cell. Instead, it learns to map control-cell distributions to perturbation-specific output distributions. SPECTRA therefore uses distributional training losses rather than paired cell-wise reconstruction losses.

### Gene regulatory network inference

SPECTRA uses an inferred gene regulatory network (GRN) as a structural prior for message passing. Numerous methods have been proposed to infer gene–gene relationships from transcriptomic and other omic data [32]. In perturbational single-cell settings, however, GRN inference should ideally recover edges that are consistent not only with observational co-expression, but also with the transcriptional consequences of targeted genetic interventions. We therefore constructed the initial parent GRN using an interventional GRN inference strategy motivated by CausalBench [33], a benchmark for network inference from single-cell perturbation data. CausalBench evaluates inferred networks using both biologically motivated reference interactions and interventional statistical criteria, including mean Wasserstein distance for predicted edges and false omission rate for omitted effects. Among the evaluated methods, the CausalBench challenge submissions Mean Difference and Guanlab were among the strongest performers, with Guanlab showing particularly strong performance on the biological evaluation [33]. We therefore used the Guanlab approach, a supervised LightGBM-based edge-ranking method developed for the CausalBench challenge [34], to obtain a high-recall parent network.

Because parent GRNs inferred from high-dimensional expression data can be dense, we further sparsified the network before using it in SPECTRA. For this step, we developed FUNGI (Functional Unraveling of Network Geometry for Inference), a dataset-agnostic graph sparsification pipeline designed to convert a dense parent GRN into a sparse topology suitable for graph neural network training. FUNGI builds on the DASH (Domain-Aware Sparsity Heuristic) [35] framework and treats edge selection as a hyperparameter optimization problem over biologically motivated topological criteria. The objective balances macro-scale properties, such as heavy-tailed degree structure, hub hierarchy, and giant-component size, with meso-scale properties, such as modularity, clustering or feed-forward-loop enrichment, and disassortative hub-to-effector wiring. The feasible ranges for these quantities are calibrated from the input expression data and parent graph, rather than from external organism-specific annotation files. The resulting graph is a sparse, modular, scale-free-like GRN that provides the structural scaffold used by SPECTRA. FUNGI details and code can be found at: ww.github.com/PatrickSheehan053/FUNGI.

### Overview of SPECTRA

To initialize node features, SPECTRA conditions gene embeddings (e.g. taken from pre-trained foundation models like scGPT [11]) on the control cell’s scalar expression profile, *x*^0^, via a FiLM-like layer [12]. For a gene *g* with embedding *e*_*g*_ ∈ ℝ ^*d*^ and expression 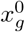, the modulated embedding *h*_*g*_ is computed as:

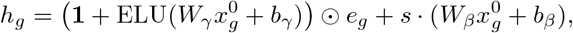

where *W*_*γ*_, *W*_*β*_ ∈ ℝ ^*d×*1^ and *b*_*γ*_, *b*_*β*_ ∈ ℝ ^*d*^ are learnable parameters, ⊙ denotes element-wise multiplication, and *s* = 0.05 scales the additive shift. Rather than using simple scalar multiplication, which uniformly scales latent dimensions and destroys the representations of unexpressed genes, this non-linear affine transformation preserves the gene’s foundational identity while dynamically adapting to its specific cellular state. The resulting modulated embeddings *h*_*g*_ are subsequently passed to the GNN, seamlessly integrating three distinct modalities: the pre-trained biological context, the cell-specific transcriptomic state, and the structural GRN topology.

The expression-conditioned node features are passed through a graph encoder composed of multiple directed graph convolution layers, defined over a given Gene Regulatory Network *G*. The encoder adopts a variational paradigm and learns a latent distribution over the basal control-cell state,

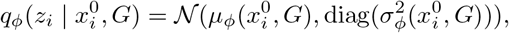

where *z*_*i*_ is a graph-structured latent representation with one latent vector per gene node. Sampling is performed through the reparameterization trick,

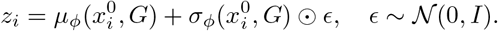

In order to construct this basal cell latent space, we use a control-cell decoder that reconstructs the unperturbed expression profile from *z*_*i*_ using the Graph Variational Autoencoder (VGAE) approach [13]. This way we ensure that the basal latent state captures the distribution of control transcriptomes before any perturbation is injected.

### Local perturbation injection

Let *g*_*p*_ be the target gene of perturbation *p*. The perturbation encoder maps the target identity into a latent perturbation vector

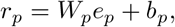

where *e*_*p*_ is the embedding corresponding to gene *g*_*p*_. Crucially, SPECTRA does not add this perturbation vector *r*_*p*_ globally to the entire cell embedding. Instead, the perturbation signal is applied only to the node corresponding to the perturbed gene:

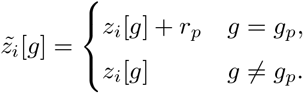

This local injection is the central modeling assumption of SPECTRA. A CRISPR intervention is treated as an initial regulatory shock at the targeted gene. The decoder must then learn how that signal propagates through the directed GRN to produce downstream transcriptomic changes.

### Directed heterophilic graph propagation

Gene Regulatory Networks (GRNs) are networks that are both directed and heterophilic with respect to transcriptomic profiles. Unlike homophilic graphs, where connected nodes tend to share similar feature values, gene regulatory edges often connect genes with divergent or opposing expression dynamics; for instance, a transcriptional repressor may downregulate its target genes producing anti-correlated behavior [32]. Conventional GNN architectures, including GCN [36], GraphSAGE [37], and GAT [38], primarily behave as low-pass filters, smoothing neighboring node representations; while effective in homophilic settings, this smoothing can erase the high-frequency signals required to model antagonistic regulatory relationships and can exacerbate over-smoothing [39].

To address this limitation, SPECTRA uses Frequency Adaptation Graph Convolutional Networks (FAGCN) [14], which adaptively combine low- and high-frequency signals through a self-gating message-passing mechanism. This allows the model to handle both homophilic and heterophilic interactions without assuming a fixed global assortativity structure. Moreover, because FAGCN learns edge-specific propagation coefficients, it can down-weight noisy or spurious regulatory links while emphasizing informative interactions, an important property given the uncertainty and false-positive rate of inferred GRNs [32].

Formally, a standard FAGCN layer updates the representation of node *i* as

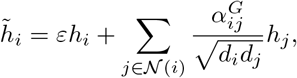

where *ε* controls the residual contribution of the original node representation, and 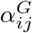 is a learned coefficient determining the relative contribution of low- and high-frequency information from node *j* to node *i*. The residual term improves training stability and mitigates over-smoothing and over-squashing [40, 41].

Since directionality in GRNs encodes regulatory flow, we adapt FAGCN to directed graphs using the Dir-GNN framework [15]. Specifically, we compute separate convolutions over incoming and outgoing edges and combine them through a weighting parameter *α*:

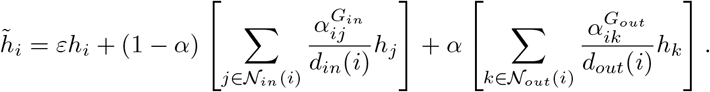

In the decoder, we set *α* = 0.7 to prioritize outgoing edges, thereby encouraging perturbation signals to propagate from the targeted gene toward its downstream regulatory targets. Degree normalization is also separated into in-degree and out-degree terms to account for the asymmetric topology of directed regulatory networks.

### Training objective

SPECTRA is trained with a hybrid objective designed for unpaired perturbation data. For control cells, the model uses a standard variational reconstruction objective:

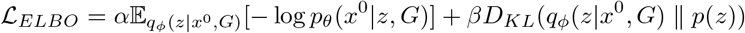

where *p*_*θ*_ is the decoder and *q*_*ϕ*_ is the encoder. This term ensures that the basal encoder learns a stable and regularized latent representation of unperturbed transcriptomes.

For perturbed cells, paired supervision is unavailable. SPECTRA therefore aligns the predicted and ground-truth perturbed distributions at the batch level using Maximum Mean Discrepancy (MMD), following the approach of STATE [8]. For a mini-batch *b* containing *S* predicted perturbed cells 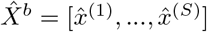 and *S* real perturbed cells *X*^*b*^ = [*x*^(1)^, …, *x*^(*S*)^], the MMD term is:

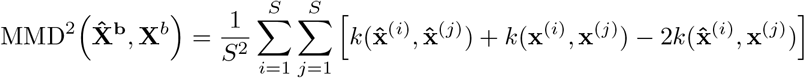

To accurately capture perturbation-specific effects, we utilize a weighted energy distance kernel for *k*(*u, v*).

While standard energy distance computes a uniform mean squared error across all *N* genes, we modulate these uniform weights 1*/N* using a differentially expressed gene (DEG) aware weighting scheme adapted from Mejia et al. [19]. Initially, DE genes are identified by comparing perturbed and control cells using a Wilcoxon rank-sum test, which yields a raw z-score (*z*_*p,g*_) quantifying the statistical significance of the perturbation effect *p* for every gene *g*. However, while z-scores effectively capture the statistical reliability of a separation, they do not account for the relative magnitude of the biological effect. In single-cell datasets with large cell populations, the Wilcoxon test can yield highly significant z-scores for biologically negligible shifts simply because the variation is consistent across hundreds of cells. Conversely, while the log-fold change (logFC) directly measures effect magnitude, it is highly sensitive to dropout noise and can produce unstable, artificially inflated values for lowly expressed genes.

To resolve this trade-off, we introduce a continuous effect-size gate that modulates the raw z-score using the absolute mean expression difference. This ensures the model prioritizes genes with both high statistical confidence and substantial biological magnitude. The composite score 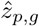 for a gene *g* under perturbation *p* is defined as:

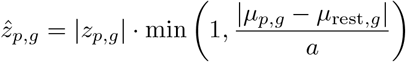

where *µ*_*p,g*_ and *µ*_rest,*g*_ represent the mean log1p-transformed expression of gene *g* in the perturbed and unperturbed (rest) populations, respectively. The hyperparameter *a* serves as a biological significance threshold. For instance, setting *a* = 0.5 ensures that any gene exhibiting a log1p expression shift of 0.5 or greater retains its full statistical z-score weight, whereas genes with shifts below this threshold are penalized linearly to suppress the influence of highly confident but biologically trivial variations.

Following the computation of the composite scores 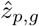, they are transformed through min-max normalization, amplified by a quadratic scaling function to widen the gap between foreground and background genes, and finally normalized to sum to one, as described in Mejia et al. [19]. This produces a strictly positive, perturbation-specific weight vector *w* = (*w*_*g*_)_*g*=1…*N*_. These final weights are then incorporated into the energy distance kernel:

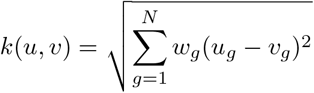

This strategy is mathematically motivated by the highly sparse nature of CRISPR perturbation data, where only a small subset of genes undergoes substantial transcriptional changes while the majority remain largely unaffected. Under conventional unweighted objectives, these biologically relevant signals can be overwhelmed by the large number of unchanged genes, encouraging models to converge toward near-mean predictions and contributing to mode collapse [19]. By explicitly prioritizing genes that are most responsive to each perturbation, this DEG-aware weighting increases sensitivity to perturbation-specific effects and improves the model’s ability to recover biologically meaningful differential expression patterns. Additionally, we introduce a Pseudobulk Cosine Similarity objective to strictly align the global direction of the transcriptomic response. The cosine similarity is defined as:

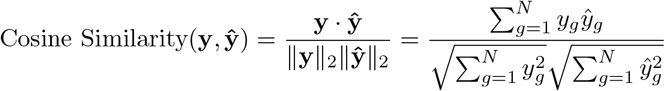

By computing the cosine similarity between the ground-truth pseudobulk expression vector 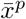 and the predicted pseudobulk vector 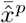 across all *N* genes, the model is penalized if the relative proportions of up- and down-regulated genes do not match the true biological signature.

Therefore, the total training objective is ℒ = *γ*ℒ_*ELBO*_ + *η*ℒ_*MMD*_ + *λ*ℒ_*cosine*_, with ℒ_*cosine*_ = 1 − cosine similarity.

### Data Preprocessing and Quality Control

Perturb-seq data were subjected to rigorous quality control and preprocessing utilizing the standardized Scanpy framework [42]. To mitigate technical noise and inherent data sparsity, genes detected in fewer than 200 cells and individual cells expressing fewer than 5 genes were strictly filtered from the raw count matrices. Following this initial filtering, library size normalization was performed by globally scaling the total unique molecular identifier (UMI) counts per cell to a target depth of 10,000. This normalization was subsequently followed by a natural logarithmic transformation to stabilize variance. To ensure sufficient statistical power for learning robust population-level distributional shifts, experimental perturbations represented by fewer than 50 viable cells were excluded from downstream analyses. We excluded perturbations targeting genes absent from the Gene Regulatory Network (GRN), as SPECTRA requires the target gene to exist as a structural node to inject and propagate the perturbation signal. Crucially, to ensure an equitable and unbiased comparative analysis, this identical preprocessing pipeline was uniformly applied to the input data of all benchmarked models.

### Pseudobulk mean baseline

As a non-parametric reference model, we implemented a pseudobulk mean baseline. This simple baseline model is designed to just summarize the average expression profile observed in the training data [31]. It preserves the marginal mean expression profile of perturbations observed during training, but it cannot model cell-to-cell variability, nonlinear perturbation effects, or perturbation-specific responses for unseen targets. In fact, for unseen perturbations, the model collapses to the global average perturbation profile, making it a stringent reference for evaluating whether SPECTRA learns information beyond pseudobulk memorization or average-expression bias.

The set of perturbations used to estimate the baseline is

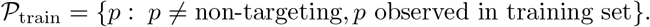

Let *x*_*i*_ ∈ ℝ^*G*^ denote the log-normalized expression vector of cell *i* over *G* genes, and let *p*_*i*_ denote its perturbation label. For each perturbation *p* observed in the training set *P*_train_ we compute a perturbation-specific pseudobulk profile

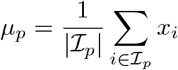

where ℐ_*p*_ is the set of training cells assigned to perturbation *p*. For perturbations not observed during training, we use a global perturbation-average profile computed across perturbation-level pseudobulks:

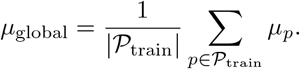

Thus, for a query perturbation *q*, the predicted mean expression vector is

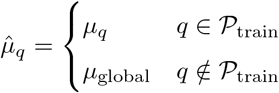

In the single-context setting considered here, this baseline is equivalent to the perturbation mean baseline used in STATE: STATE defines the perturbation mean as a control-referenced perturbation offset averaged across training contexts and added to the target-context control mean [8]. When only one cellular context is present, this expression simplifies to the perturbation-specific pseudobulk mean itself.

### Evaluation metrics

Recent research highlights that a fundamental challenge in the field is designing a common, biologically meaningful evaluation framework [19]. Traditionally, the predictive performance of these models has been assessed using global, alignment-based metrics such as Mean Squared Error (MSE), Mean Absolute Error (MAE), Pearson correlation, and coefficient of determination (*R*^2^). While these metrics provide a mathematical summary of overall transcriptomic similarity across thousands of genes, they are fundamentally misaligned with the biological reality of cellular perturbation responses [19, 31]. The underlying issue is that transcriptomic responses to genetic perturbations are inherently sparse, typically modulating only a small highly specific subset of the genome. Because the vast majority of genes remain unaffected during a given intervention, models are susceptible to learning “lazy” representations, achieving deceptively high scores on global metrics simply by predicting near-control expression levels or by capturing the average perturbation effect across the dataset. Recent benchmarking studies, such as the Systema framework [20], demonstrate that relying on these traditional metrics often rewards models for capturing systematic background variation rather than true, perturbation-specific causal signals. Ultimately, we think the practical biological utility of perturbation forecasting relies not on the minimization of global expression error, but on the precise identification of Differentially Expressed Genes (DEGs). Therefore, evaluation paradigms must consider not only standard regression metrics but also metrics quantifying DEG overlap, which directly assess the proportion of accurately predicted DEGs relative to empirical post-perturbation profiles.

To this end, we adopt the evaluation suite proposed in recent benchmarking literature [16], which encompasses both population-average and population-distribution metrics such as Mean Squared Error (MSE), E-distance, the Pearson Correlation Coefficient of the delta (PCC-delta), Wasserstein distance, and Kullback-Leibler (KL) divergence. To accurately assess a model’s capacity to capture perturbation-specific causal signals, these metrics can be computed exclusively on the top *N* groundtruth DEGs.

To rigorously quantify DEG overlap, we formulate DEG prediction as a binary classification task: determining whether a specific gene is significantly differentially expressed in response to a given perturbation. This framing enables the application of classical binary classification metrics. However, in binary classification, simultaneously minimizing false positives (FPs) and false negatives (FNs) is mathematically impossible, and an optimal trade-off must be established. For the practical application of *in silico* perturbation forecasting, we argue that minimizing FPs (i.e., maximizing precision) is of paramount biological importance. If a virtual cell model identifies a gene as a DEG, there must be high statistical confidence in that prediction to justify the allocation of resources for subsequent *in vitro* experimental validation. Consequently, we emphasize the precision score

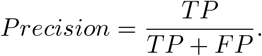

which directly quantifies the probability that a predicted DEG is a true positive. Additionally, we report the F1 score to provide a more balanced metric combining precision and recall:

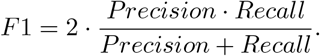

For these hard-classification metrics, for both the ground truth data and the generated data genes were classified as DEGs using a Wilcoxon rank-sum test with False Discovery Rate (FDR) correction (*α* = 0.01) and an absolute log-fold change (| log_2_ FC|) threshold of 0.3.

It is critical to acknowledge, however, that empirical DEG inference is an inherently noisy process. The resulting gene sets are highly sensitive to the chosen statistical methodology [43], as well as the strictness of the applied p-value and logFC thresholds. Consequently, evaluating DEG overlap strictly as a hard-classification problem can introduce misleading boundary artifacts; for instance, given a strict | log_2_ FC| threshold of 0.3, a gene exhibiting a predicted | log_2_ FC| of 0.29 against a ground-truth | log_2_ FC| of 0.31 would be strictly penalized as a false negative, despite the model capturing the correct biological trend.

To mitigate the limitations of threshold-dependent hard classification, we incorporate the Area Under the Precision-Recall Curve (AUPRC) metric proposed in [18]. By treating the predicted expression shift as a continuous confidence score and calculating precision and recall across all possible threshold values, the AUPRC provides a robust, threshold-agnostic evaluation of the model’s ability to rank and identify true DEGs. We implemented this evaluation pipeline in Python, utilizing the Wilcoxon rank-sum test as the core statistical framework.

